# Addition of Degenerate Bases to DNA-based Data Storage for Increased Information Capacity

**DOI:** 10.1101/367052

**Authors:** Yeongjae Choi, Taehoon Ryu, Amos C. Lee, Hansol Choi, Hansaem Lee, Jaejun Park, Suk-Heung Song, Seoju Kim, Hyeli Kim, Wook Park, Sunghoon Kwon

## Abstract

DNA-based data storage has emerged as a promising method to satisfy the exponentially increasing demand for information storage. However, practical implementation of DNA-based data storage remains a challenge because of the high cost of DNA per unit data. Here, we propose the use of eleven degenerate bases as encoding characters in addition to A, C, G, and T, which increases the information capacity (the amount of data that can be stored per length of DNA sequence designed) and reduce the cost of DNA per unit data. Using the proposed method, we experimentally achieved an information capacity of 3.37 bits/character, which is more than twice when compared to the highest information capacity previously achieved. Finally, the platform was projected to reduce the cost of DNA-based data storage by 50%.

## Main Text

The annual demand for digital data storage is expected to surpass the supply of silicon in 2040, assuming that all data are stored in flash memory for instant access^1^. Considering the massive accumulation of digital data, the development of alternative storage methods is essential. One alternative is DNA-based data storage, which converts the binary digital data of 0 and 1 into the quaternary encoding nucleotides A, C, G, and T, synthesizes the sequence, and stores the data^2^. This concept^2-9^ is attractive due to two main advantages: the high physical information density of petabytes of data per gram and the durability, as the storage lasts for centuries without energy input. Due to these advantages, DNA-based data storage is expected to supplement the increasing demand for digital data storage, especially for archival data that are not frequently accessed. Since DNA-based data storage was proposed, the major goal was to improve data encoding algorithms to reduce data error or loss considering the biochemical properties while handling DNA. For example, algorithms have been proposed to remove high GC content and long homopolymers during encoding, which are known to cause errors^8,9^. In addition, various error correction algorithms for DNA-based data storage^3,5,6,8,9^ have been developed to correct errors or recover dropped data fragments during DNA amplification and sequencing. These previous studies on encoding algorithms showed 100% reconstruction of the data from DNA.

The next step towards the practical use of DNA-based data storage is to reduce the cost of storing the data. The cost of DNA-based data storage is categorized into the cost of data writing through DNA synthesis and the cost of data reading through DNA sequencing. Among these two costs, the cost of data writing is predominant because it is tens of thousands times more expensive per unit nucleotide than that of reading. However, previous studies have shown that DNA can be put to practical use as a backup storage medium only when the cost of the data writing is approximately 100 times less^3^. Therefore, it is essential to minimize the amount of data that can be stored per length of DNA sequence that is designed (information capacity, bits/character). Previous methods have a theoretical information capacity limit of *log*_2_*4*, or 2.0 bits/character, because DNA comprises four encoding characters (A, C, G, T) (Supplementary Note 1). If additional encoding characters are introduced, the information capacity of *log*_2_(number of encoding characters) dramatically increases, further reducing the cost of DNA data storage.

Here, we propose and demonstrate the use of degenerate bases (combination of the four DNA bases that can be inserted at any base sites within a sequence)^10^ as additional encoding characters to exceed the theoretical information capacity limit of 2.0 bits/character. Degenerate bases are located in the DNA sequence when nucleotides are mixed at a specific position in the DNA sequence. For example, in the sequence ‘AWC’, ‘W’ indicates a combination of A and T; thus, two types of nucleotide variants exist in the pool of molecules: ‘AAC’ and ‘ATC’. Ideally, synthesizing oligonucleotides with degenerate bases does not require additional cost during the synthesis for data writing. Even if the number of data reads in sequencing is increased to identify the variants, the additional cost is still less than 5% of the total data writing cost. In this letter, by using all eleven degenerate bases in addition to the four DNA characters, we experimentally achieved an information capacity of 3.37 bits/character and shortened the DNA length needed to store the same amount of data by more than half compared to previous reports^2–5,8,9^. Finally, the proposed platform is projected to reduce the cost of DNA-based data storage by 50% by reducing the data writing cost and increasing the data reading cost.

The conversion from a four to a fifteen character-based encoding system theoretically allows a maximum information capacity of 3.90 bits/character (*log*_2_15) (previously 2.0 bits/character, (*log*_2_4)) and shortens the length of DNA required to store an equivalent amount of data by approximately half (Fig. 1a). While previous research has increased the information capacity to near the 2.0 bits/character theoretical limit by optimizing the data to DNA encoding algorithm, our approach increases the information capacity by increasing the theoretical limit (Fig. 1b). The degenerative portion of the encoded sequence, which can increase the theoretical limit, is incorporated by mixing the DNA phosphoramidites during the synthetic procedure^11^ and generating variants of the corresponding combinations of A, C, G and T (Fig, 1C, D). Ideally, for column-based^11^ and inkjet-based^12–14^ oligonucleotide synthesis, degenerate bases can be added without extra cost because the total amount of phosphoramidites used is the same(Supplementary Note 2). Also, current synthesis techniques synthesize more than billion molecule of oligonucleotides molecules per design, which are sufficient to generate variant pool for degenerate base. Therefore, the platform shortens the length of DNA to store the equivalent amount of data by approximately half, decreasing the expense for DNA synthesis, i.e., the data writing.

**Figure 1.**
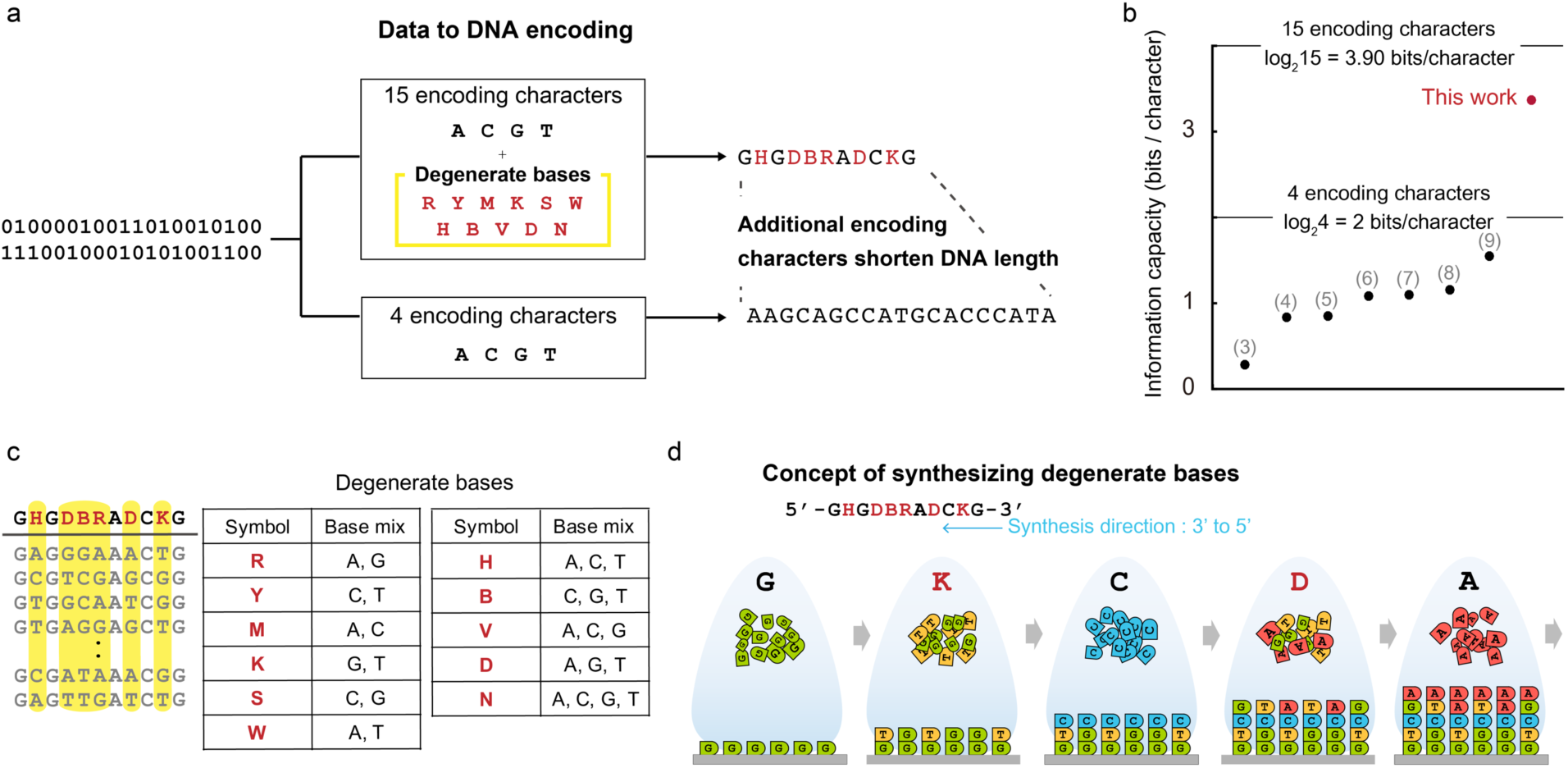
DNA-based data storage with added degenerate bases enables increased information capacity. (**a**) Binary data is encoded to DNA sequences comprising not only the 4 traditional encoding characters A, C, G, and T but also 11 additional degenerate bases. The length of encoded DNA is less than that of the four-character encoding method. (**b**) The theoretical information capacity limit is therefore increased from 2 bits/character to 3.9 bits/character. The dots in the graph describe the information capacity values in previous research, and the numbers indicate the corresponding reference. (**c**) A degenerate base represented by an encoding character describes a mixed pool of more than two types of nucleotides. (**d**) Degenerate bases can be generated by mixing the DNA phosphoramidites during the synthesis.

We encoded an 854 bytes-text file to DNA sequences (Fig. 2, Supplementary Fig. 1). The data were transformed into a series of three-character DNA codons, the sequence of which consists of three encoding characters. The last base in the sequence of the codons was designed to not be equivalent to the front-most base in the sequence of the next codon to avoid the generation of homopolymers of 4 nt or more (Supplementary Table 1). The encoded information was divided into 42 nt fragments, and an address composed of 3 nt of non-degenerate bases (Supplementary Table 2) was assigned to each fragment (Fig. 2a). Each fragment was supplemented with two adapters (20 nt each for the 5’ and 3’ end) for amplification and sequencing, and the entire fragment was 85 nt in length. From the design described, 45 DNA fragments were synthesized by the column-based oligonucleotide synthesizer without additional cost. Considering the number of bits encoded in the total nucleotide synthesis excluding the adapters, an information capacity of 3.37 bits/character was achieved experimentally, which is more than twice the highest reported value of 1.57 bits/character^9^. The information capacity demonstrated was lower than the theoretical maximum because the encoding efficiency was lowered to avoid homopolymer sequences and incorporate non-data address sequences for each fragment. The synthesized DNA library consisting of approximately 800 molecules was amplified by designed adapters and was sequenced by an Illumina MiniSeq. The raw data was filtered using the designed length and categorized by addresses. Then, the duplicated reads were removed and the distribution of A, C, G, and T in each position on the fragment was analyzed (Fig. 2b). The intermediate ratio of the nucleotides analyzed was not consistently equivalent because the coupling efficiency during synthesis varies for each base, by type and position in the growing oligonucleotide^15,16^. However, when we observed the ratio of A:C:G:T in the sequence analyzed at the same position using a scatter plot, the points were clustered into fifteen groups, eleven of which had an intermediate ratio of more than two bases considered degenerate bases (Fig. 2c). The other four that had a dominant ratio of a particular nucleotide were considered pure sequences. Through this decoding process, we successfully recovered the original data from the raw next-generation sequencing (NGS) data. We also recovered the data in 10 of 10 cases when randomly down-sampled to the average coverage of 250x. If the average NGS coverage is lower than 250x, the error rate increases because the intersections between the clusters of encoding characters are augmented. To demonstrate the scalability of the introduced platform, we also stored 135.4 kB of data (Supplementary Fig. 2) in 4503 fragments of DNA using the pooled oligonucleotide synthesis method, which is cost effective and high throughput. To manage the error^17^ and amplification bias that may occur when synthesizing and amplifying oligonucleotide pools with high complexity^18,19^, we added Reed-Solomon-based redundancy^8^ (Supplementary Note 3, Supplementary Fig. 3). Even though only two degenerate bases, W and S, were used for this demonstration due to equipment constraints (Supplementary Note 4), an information capacity of 2.0 bits/character was achieved. We recovered the data in 10 of 10 cases when randomly down-sampling the average coverage to 250x. This is higher than the minimum NGS coverage required for DNA-based data storage without degenerate bases, which is approximately 5x^7^. We summarized our experimental results in terms of the input data, number of oligonucleotides, minimum coverage, physical density, and information capacity (Fig. 2d). Physical density describes relation between molecule number used and data quantity, while information capacity describes that between designed character number and data quantity. Although we synthesized oligonucleotide variants in single designed fragments to incorporate the degenerate bases, fewer oligonucleotide molecules per fragment (hundreds) were sufficient to decode the data, than that in a previous report^9^. In this respect, we renewed the highest experimentally proven information capacity and physical density by compromising higher NGS coverage.

**Figure 2.**
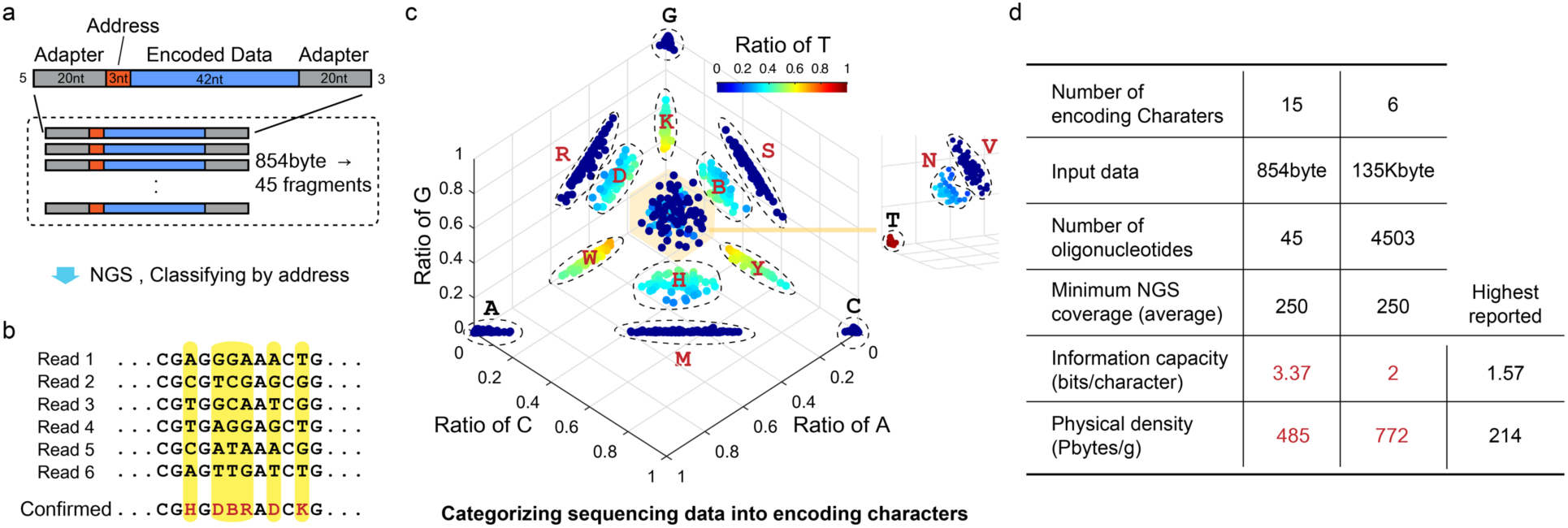
Structure and decoding result of the DNA-based data storage platform. We achieved the highest information capacity and physical density of DNA-based data storage. (**a**) Design structure of DNA fragments. (**b**) DNA fragments can be analyzed using NGS. After classification by address, degenerate bases can be decoded by examining the distribution of characters in the same position (yellow bar). (**c**) Degenerate bases can be determined from the scatter plot of the ratio of bases in the same position. (**d**) Summary of the experimental results. The information capacity is calculated from the input information in bits divided by the number of encoding characters (excluding that of adapter sites). We compared the results of our work with those of Erlich and Zielinski *(9)*, who previously reported the highest information capacity and physical density using pooled oligo synthesis and high-throughput sequencing data. The physical density is the ratio of the number of bytes encoded to the weight of the DNA library used to decode the information.

In addition to the experimental results, we simulated the error rate of the platform in terms of NGS coverage for data recovery when various types of degenerate bases are used on a large scale. Because the call frequency of each base comprising the degenerate bases follows a binomial distribution (Supplementary Fig. 5, Supplementary Note 5), the platform was modeled using Monte Carlo simulation. We simulated the error rate per base pair of the models by using various sets of degenerate bases (Fig. 3a) when fragments are represented unevenly due to amplification bias (Supplementary Figure 6). The assumed length of the fragment used in the simulation was 200 nt with a 20-nt adaptor at both ends, and the data was stored at 148 nt, except for the address of 12 nt. In the simulation, we also introduced additional characters specified by two nucleotides with different ratios (e.g., W1 for A:T=3:7 and W2 for A:T=7:3) and expanded the number of encoding characters to 21. The data show that the use of various types of degenerate bases increases the error rate but the error rate decreases with increasing NGS coverage. Given NGS coverage of 1300x or more, decoding 100 MB with 10% Reed-Solomon redundancy in all proposed cases can proceed without error. As a result, we achieved 2.67 bits/character when using 15 encoding characters and 3.05 bits/character when using 21 encoding characters. Although the platform requires high NGS coverage, the sequencing technology has a rapid speed of evolution, and the current DNA sequencing cost per base is approximately 50,000 times lower than the synthesis cost per base used for DNA-based data storage^3,9,20^. Moreover, since cost of DNA sequencing is decreasing faster than the Moore’s law and faster than that of DNA synthesis, the price gap between the sequencing and synthesis will increase by orders, if the current trend continues^1,21^. Even if the proposed platform has 2000x NGS coverage as an extreme case, the data reading cost will be less than 5% of the writing cost and less than 0.5%, which will be ignorable, in five years (Fig. 3b). Finally, when the pool-based oligonucleotide synthesizer is set for degenerate base synthesis, the proposed platform is projected to reduce the cost of DNA-based data storage to $2052/1MB when using 15 encoding characters and $1795/1MB when using 21 encoding characters, which is approximately 50% of the previous minimum of $3555/1MB ^9^ (Fig. 3b, Supplementary Note 6).

**Figure 3.**
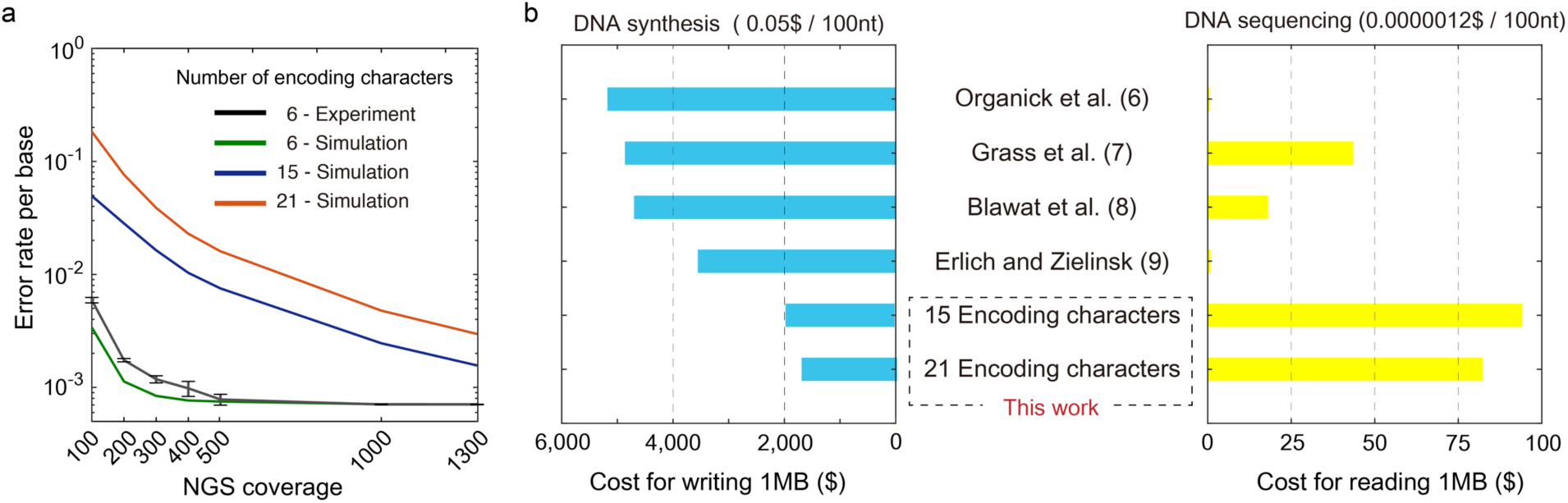
The proposed platform is projected to reduce the cost of DNA-based data storage by 50%. (**a**) The error rate per base pair according to the average NGS coverage over all fragments. The black line shows the experimental results, and the other three lines represent the Monte Carlo simulation results. For the experiment and simulation shown in green, we used A, C, G, T, W, and S for encoding. For the simulation shown in blue, we used A, C, G, T and all other degenerate bases. For the simulation shown in red, we used A, C, G, T, [R, Y, M, K, S, W – ratio of bases mixed of 3:7 and 7:3], H, V, D and N. The standard deviation (s.d.) of the experimental results were obtained by repeating the random sampling 5 times. The error bars represent the s.d. (**b**) Cost projection of previous research and the proposed platform. We used the cost of column-based DNA synthesis reported by Erlich and Zielinski^9^. The cost of DNA sequencing was reported by K. Wetterstrand ^20^. We used A, C, G, T and all other eleven degenerate bases as encoding characters. Additionally, we used A, C, G, T, [R, Y, M, K, S, W – ratio of bases mixed of 3:7 and 7:3], H, V, D and N as 21 encoding characters. The numbers indicate the corresponding reference. Details on the projection method are described in the Supplementary Note 6.

In this demonstration, by utilizing degenerate bases, the information capacity and physical density were more than doubled compared to those of previously reported DNA-based data storage systems. In particular, as the information capacity increases, the platform shortens the length of DNA required to store an equivalent amount of data and could possibly decrease the total expense of data storage by half. To realize what is simulated in this study, future development in oligonucleotide synthesis will be necessary. First, an oligonucleotide pool synthesis setup can be used to increase the information capacity by incorporating all the degenerate bases in the encoding characters by addition of the nozzles. Second, if the synthesis setup can precisely control the ratio of the nucleotides consisting degenerate base is developed, even more encoding characters can be used. Currently, no method that precisely controls the ratio has been reported in our knowledge and this could not be done by simply changing the ratio of input phosphoramidite, as shown in our experiments and in the literature in the past^15,16^. If, with future research, it is possible to optimize the platform for a large-scale experiment and to generate modified degenerate bases with non-equivalent ratios suggested in the simulation. Additionally, if synthesis and sequencing methods for synthetic bases^22^ are developed, they can be used as another type of encoding characters. In addition to the development of these synthetic methods, reduction in the DNA amplification bias will improve the practical efficiency of the method. Together with these additional technologies, the proposed platform with increased information capacity will enable the practical use of the DNA-based data storage in the future.

## Acknowledgments

This work was supported by Samsung Research Funding Center of Samsung Electronics under Project Number SRFC-IT1601-08. Authors acknowledge gratitude towards Duhee Bang (Department of Chemistry, Yonsei University, Seoul) for productive discussion on manuscript.

## Author contributions

Yeongjae Choi, Taehoon Ryu, Wook Park and Sunghoon Kwon initiated and designed the experiments. Yeongjae Choi, Amos C. Lee, Wook Park and Sunghoon Kwon wrote the manuscript. Yeongjae Choi, Taehoon Ryu, Amos C. Lee, Hansol Choi, Hansaem Lee, Jaejun Park, Suk-Heung Song, Seoju Kim, and Hyeli Kim conducted the research including DNA synthesis and analysis.

## Supplementary Materials

Methods

Figures S1-S6

Tables S1-S4

